# Conformational Properties of Polymers at Droplet Interfaces as Model Systems for Disordered Proteins

**DOI:** 10.1101/2023.07.29.551102

**Authors:** Jiahui Wang, Dinesh Sundaravadivelu Devarajan, Arash Nikoubashman, Jeetain Mittal

**Affiliations:** Artie McFerrin Department of Chemical Engineering, Texas A&M University, College Station, TX 77843, United States; Institute of Physics, Johannes Gutenberg University Mainz, Staudingerweg 7, 55128 Mainz, Germany; Department of Chemistry, Texas A&M University, College Station, TX 77843, United States; Interdisciplinary Graduate Program in Genetics and Genomics, Texas A&M University, College Station, TX 77843, United States

**Keywords:** Liquid–liquid phase separation, membraneless organelles, interfacial behavior, molecular simulations

## Abstract

Polymer models serve as useful tools for studying the formation and physical properties of biomolecular condensates. In recent years, the interface dividing the dense and dilute phases of condensates has been discovered to be closely related to their functionality, but the conformational preferences of the constituent proteins remain unclear. To elucidate this, we perform molecular simulations of a droplet formed by liquid–liquid phase separation of homopolymers, as a surrogate model for the prion-like low-complexity domains. By systematically analyzing the polymer conformations at different locations in the droplet, we find that the chains become compact at the droplet interface compared to the droplet interior. Further, segmental analysis revealed that the end sections of the chains are enriched at the interface to maximize conformational entropy, and are more expanded than the middle sections of the chains. We find that the majority of chain segments lie tangential to the droplet surface and only the chain ends tend to align perpendicular to the interface. These trends also hold for the natural proteins FUC LC and LAF-1 RGG, which exhibit more compact chain conformations at the interface compared with the droplet interior. Our findings provide important insights into the interfacial properties of biomolecular condensates and highlight the value of using simple polymer physics models to understand the underlying mechanisms.

---

Membraneless organelles or biomolecular condensates formed through liquid*–*liquid phase separation (LLPS) have been widely reported in various cellular functions, including gene expression, signal transduction, stress response, and the assembly of macromolecular complexes^1–5^. Examples of such condensates include the nucleolus, Cajal bodies, P bodies, and stress granules. Intrinsically disordered proteins (IDPs) play an important role in the formation of biomolecular condensates through LLPS^6–11^. Due to the numerous similarities between IDPs and synthetic polymers, classical polymer-based models offer a powerful approach for investigating the conformations^12^, dynamics^13^, and phase behavior of IDPs^14–17^. In particular, such models have been extensively used to reveal the sequence-dependent conformations of IDPs in the dense and dilute phases of condensates^18–27^.

In recent years, there has been increasing recognition of the important role played by the interfaces dividing the dense and dilute phases of biomolecular condensates. Folkmann *et al*. have reported that the IDP assemblies at the interface of PGL-3 droplets reduced the surface tension, thus preventing droplet coarsening. Kelley *et al*. reported that amphiphilic proteins on the surface of condensates acted like surfactants, thus regulating the size and structure of condensates^28^. Further, the interface can affect the interactions between biomolecular condensates and other biomolecules within the cell^29^. Böddeker *et al*. demonstrated that cytoskeletal filaments had a nonspecific affinity for stress granule interfaces, which explained the distinct enhancement of tubulin density around granules^30^. Lipiński *et al*. reported that the condensate interface can serve as a nucleation site promoting the aggregation of amyloidogenic proteins^31^. Inspired by the critical role of interfaces in various functionalities, much research has focused on determining mesoscopic interfacial properties such as surface tension^13,32,33^, surface adsorption^34^, and electrochemical properties^35^, which are deemed important for regulating biological functions^29,36^. For example, it was shown that the condensates could form an autophagosome only if the surface tension is lower than a critical value^37^. The film formed by protein adsorption at the air/water interface has a low interfacial elasticity at the protein’s isoelectric pH because of the compact structure^38^.

However, the microscopic conformations of individual IDPs at interfaces remain largely unclear, even though these exposed polymers largely dictate the surface properties. . For example, surfaces grafted by a synthetic polymer such as PNIPAM have been shown to result in a four-fold difference in the Young’s modulus between the swollen and collapsed states,^39^ and the surface wettability is stronger for polymer interfaces in stretched states than in collapsed states^40^. For biomolecular condensates, the chain conformations can be important for their functions like the regulation of condensates conformation on mRNA decapping^41^. To shed light on this question, Farag *et al*. recently studied the interface conformations of biomolecular condensates using lattice-based Monte Carlo simulations^42,43^.They reported that the overall dimensions of prion-like low-complexity domains and homopolymers varied within the condensates, being most *expanded* at the interface and preferring to be oriented *perpendicular* to it. These pioneering studies open up avenues for detailed characterization of the segmental-level conformational properties and their contribution to the chain-level conformational preferences at the condensate interfaces, which remain elusive.

In this letter, we report chain-level and segmental-level conformations of IDPs at the condensate interface from molecular simulations of hydrophobic homopolymers and two naturally occurring IDPs. We use an off-lattice polymer model, where each residue (“monomer”) is represented by a spherical bead in an implicit solvent (model details are provided as Supporting Information (SI)). This model is a good approximation for numerous IDPs, such as prion-like low-complexity domains^42^. Recently, we successfully used hydrophobic homopolymers as a reference to establish the biophysics of phase separation of IDPs, revealing that distributed interactions better stabilize the condensed phase than localized interactions^24^. To understand the basic principles governing the chain conformations in the droplet, we focus here on the hydrophobic homopolymers and include the results for the natural IDPs in the SI.

Due to the hydrophobic nature of the employed model IDP, it quickly formed a spherical condensate of radius *R* = 249Å, which is defined as the position where the concentration drops to half the value near the droplet center (**Fig.** 1). Drawing upon previous research on the liquid-vapor interface in slabs^44^, the radial concentration profile is fit to a hyperbolic tangent functional form

**Fig. 1.**
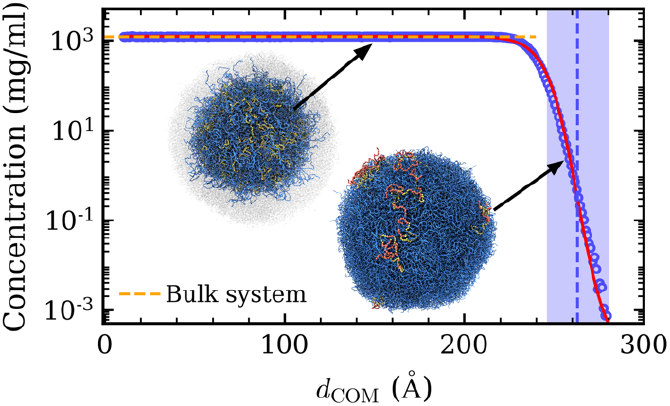
Concentration as a function of distance from the droplet’s center of mass (*d*_*COM*_). The horizontal dashed line represents the concentration in the bulk system. The red line is the fitted curve for the simulation data (symbols). The purple shaded area is the interface region. The vertical dashed line represents the middle of the interface. The inset shows simulation snapshots of the droplet interior and interface. The yellow chains in the interior snapshot are at a distance of *d*_*COM*_ = 150 Å. The chains colored yellow and red in the interface snapshot have their center-of-mass in the interface, with the red beads representing segments in the interface.

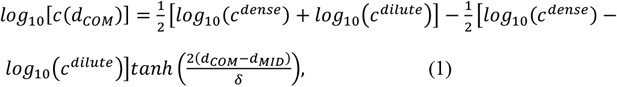

where *d*_*COM*_ is the distance from the droplet’s center of mass, *c*(*d*_*COM*_) is the radial monomer concentration, *c*^*dense*^ and *c*^*dilute*^ are the monomer concentrations of the dense phase and dilute phase, respectively, *d*_*MID*_ is the midpoint of the interface, and *δ* is the width of the interface. By fitting the computed radial concentration curve to Eq. (1), the interfacial region was defined (**Fig.** 1). To establish a bulk reference without interface effects, we simulated a concentrated homopolymer solution in a cubic box at a constant pressure of *P* = 0 atm. With increasing *d*_*COM*_, the concentration in the droplet interior remained constant (equal to the expected concentration in the bulk system), while it monotonically decreased at the interface. In this case, the dilute phase concentration is equal to 0 *mg*/*ml*, as all chains are inside the droplet.

After determining the droplet interface, we studied the conformations of polymer chains within the interface and dense phase of the droplet. For this purpose, we computed the radius of gyration of the chains^45^

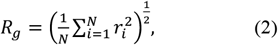

where the sum goes over the *N* monomers in the chain, and *r*_*i*_ is the distance from the *i*^th^ monomer to the chain’s center of mass (COM). We find that the corresponding *R*_*g*_ distribution in the bulk system (**Fig.** 2) was significantly broader, as compared to that of the single chain (**Fig. S**1), highlighting the preference of chains to expand inside the condensate, as expected from Flory’s ideality hypothesis^46^. Next, we analyzed the distribution of *R*_*g*_ at different positions within the droplet. Based on the distance between the chain’s COM and the droplet’s COM (*d*_*COM*_), we assigned the polymer chains into different positional bins (**Fig.** 2**a**). We found that when *d*_*COM*_ ≤ 210 Å, where the local monomer concentration is equal to the bulk system concentration (**Fig.** 1), the *R*_*g*_ distributions overlapped with the bulk distribution, indicating the consistency between the interior of the droplet and the bulk phase. With increasing *d*_*COM*_, the peak of the *R*_*g*_ distribution shifted gradually to smaller *R*_*g*_ values (**Fig.** 2**a**), demonstrating a continuous transition from expanded conformations to more compact conformations when moving from the interior of the droplet towards the interface. This result is also supported by the average number of interchain and intrachain contacts for each monomer (**Fig.** 2**b**), which decreased monotonically with increasing *d*_*COM*_, because of the decreasing monomer concentration (**Fig.** 1). At the interface, the number of average intrachain contacts was larger than the number of interchain contacts, which means that the chains interacted primarily with themselves and thus were more compact than in the droplet interior, where interchain interactions dominated.

**Fig. 2.**
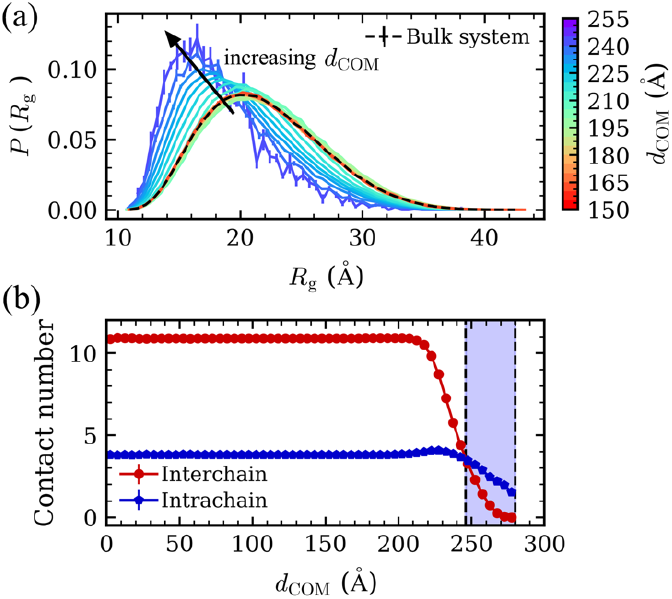
(a) Distribution of radius of gyration (*R*_*g*_). The dashed line represents the *R*_*g*_ distribution in a bulk system. The color gradient from red to purple corresponds to the distribution of *R*_*g*_ for chains with *d*_*COM*_ ranging from 150 to 255 Å. (b) Average contact number per monomer as a function of *d*_*COM*_ for the interchain and intrachain contacts. The vertical dashed lines represent the interface boundaries.

Our observation of compact chain-level conformations at the interface led us to investigate how segments within a chain contributed to its compaction, following previous calculations for the end-to-end vector of grafted polymers^47^ and *R*_*g*_ of polymer thin films^48^. We first characterized the distribution of chain segments of different length *n*_*seg*_ at the droplet interface (**Fig.** 3**a**,**b**). The middle bead of the segments for a given *n*_*seg*_ serves as the reference bead, the index of which we refer to as segment index. Note that a segment index closer to 1 or 50 corresponds to a segment closer to the chain termini, and the curves are symmetrical because the homopolymer chains do not have any “heads” or “tails”. We found that the highest probability always occurred at the position closest to the chain’s ends at the interface, regardless of the length of the segment, suggesting that the chain terminals were more inclined to distribute at the interface compared to the middle of the chains. This is in stark contrast to the droplet interior, where we found the same probabilities for segments at different relative positions within the chain (**Fig.** 3**b**). The enrichment of chain ends at the interface can be understood by considering the loss in conformational entropy incurred by placing a monomer at the interface, which is smaller for the end monomers compared to any other monomer.

**Fig. 3.**
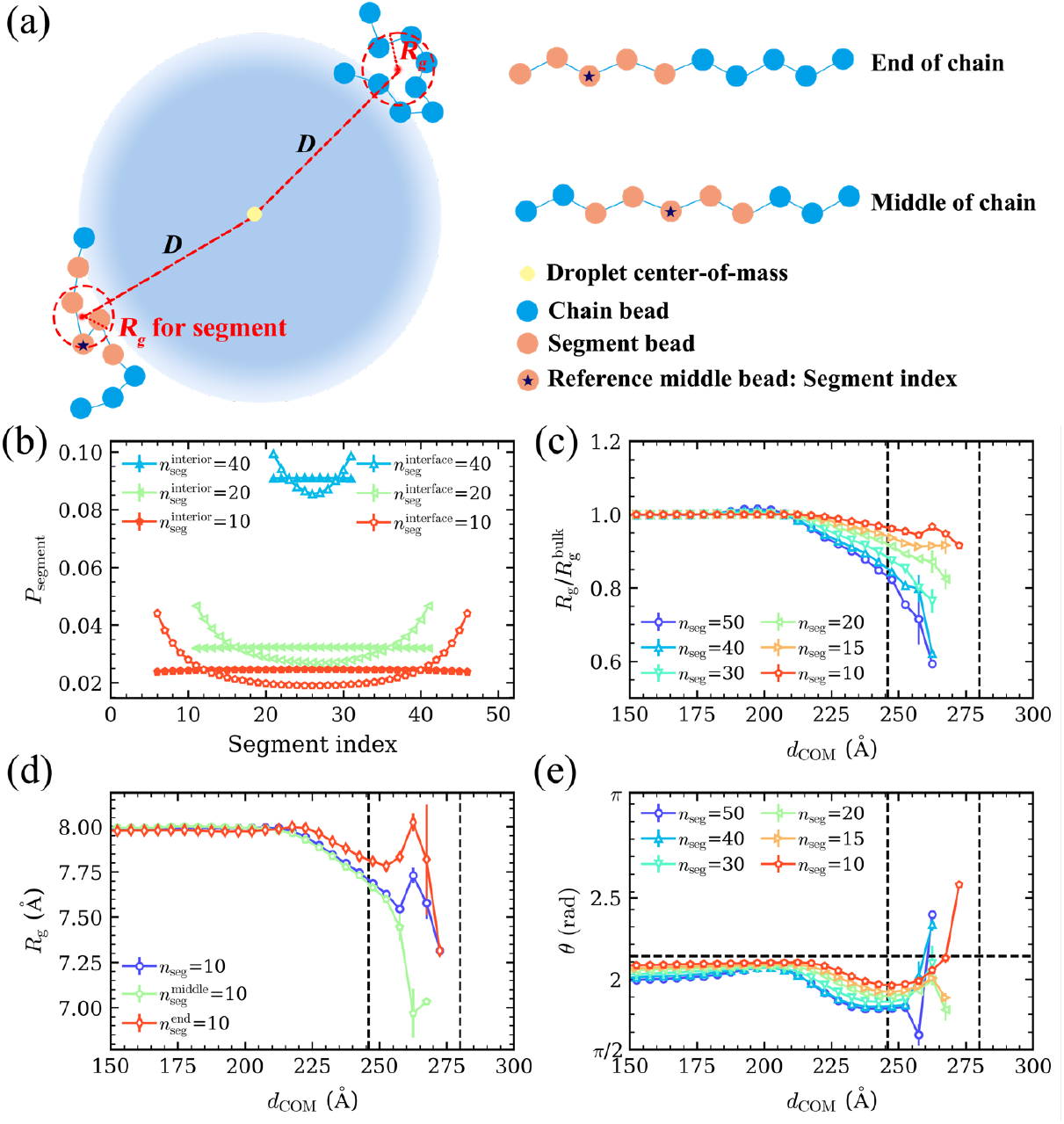
Segmental analysis. (a) Schematic diagram of the *R*_*g*_ calculation method for chains and their segments. Blue beads represent the chain monomers, while orange beads represent a specific segment. (b) Probability distribution of the segments for different segment lengths (*n*_*seg*_) at the interface (hollow symbols) and in the droplet interior (solid symbols). The segment index is represented by the index of middle bead of each segment (reference bead). When *n*_*seg*_ is even, the larger index between the two middle beads is considered as the segment index. (c) Normalized average *R*_*g*_ of the different length segments with respect to *d*_*COM*_. The black dashed lines represent the boundaries of the interface. (d) *R*_*g*_ of segments consisting of 10 monomers as a function of *d*_*COM*_. The purple line represents for the average *R*_*g*_ for all segments in the chain of that length. The green and red lines represent the segments located at the middle and end of the chains, respectively. (e) Angle (*θ*) of the different length segments with respect to *d*_*COM*_. The horizontal dashed line represents the average angle for isotropic distribution of segments.

We next asked if the spatial preference of the chain segments at the droplet interface *vs* its interior had an effect on the segmental-level conformations. For this purpose, we calculated the average *R*_*g*_ for the segments of different lengths *n*_*seg*_ as a function of distance *d*_*COM*_ between their COM and the droplet’s COM (**Fig.** 3**c**). We found that the *R*_*g*_ at the interface is smaller than that in the droplet interior for all *n*_*seg*_ considered. For *n*_*seg*_ > 15, *R*_*g*_ monotonically decreased as the droplet interface is approached. For *n*_*seg*_ = 15, *R*_*g*_ no longer decreased in the interface region. Interestingly, for *n*_*seg*_ = 10, we found that *R*_*g*_ marginally increased in the middle of the interface region, following which it decreased again. To explain this nonmonotonic trend in *R*_*g*_ for the short segments at the interface, we analyzed *R*_*g*_ of different *n*_*seg*_ located only at the chain ends and compared it with that in the middle of the chain. For longer segments *n*_*seg*_ ≥ 20, the segmental *R*_*g*_ monotonically decreased from the droplet interior to the interface, regardless of the position of the segments (**Fig. S**2). However, for shorter segments *n*_*seg*_ ≤ 15 (Figs. 3**d** **and S2**), we found that the *R*_*g*_ of the segments located at the chain ends exhibited a nonmonotonic trend in the interface region: we observed an increase in segmental *R*_*g*_ for *d*_*COM*_ > 250 Å, attaining the maximum value in the middle of the interface region, following which it continued to decrease. Surprisingly, the *R*_*g*_ of shorter segments located in the middle of the chains monotonically decreased throughout the interface region. These observations indicate that the chain termini are the only contributors to the observed expansion of short segments (**Fig.** 3**c**). In summary, although the *whole* chain is more compact at the interface, short terminal segments are more expanded than middle segments at the interface and even slightly more expanded than the short segments in the droplet interior.

Given the stark differences in the global and local conformations of the chains in the interface region, we next investigated their chain-level and segmental-level orientations. We defined the angle *θ* between the segment-to-droplet COM vector and the segment end-to-end vector to characterize the orientation of the chains in the interface region (**Fig. S**3**a**). The positions of segments are again defined through *d*_*COM*_. When the angle is close to *π*/2, the segment lies tangential to the droplet surface, whereas an angle close to *π* corresponds to an orientation perpendicular to the droplet surface. For different segment lengths (*n*_*seg*_ = 10, 15, 20, 30, 40) and the whole chain (*n*_*seg*_ = 50), the angles show an increasing trend at the interface (**Fig.** 3**e**). This increase primarily originates from the segments near the chain ends (**Fig. S** 3**b**) rather than segments in the middle of the chains (**Fig. S**3**c**). This observation can be explained by considering the end-to-end distance (*R*_*e*_) (**Fig. S**4). The smallest angle for segments (**Fig. S**3**a**) can be estimated as:

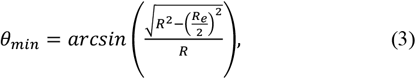

where *R* is the distance between the terminal bead of the segment and the droplet COM, and *R*_*e*_ is the end-to-end distance of the segment. At the interface, *R* is much larger than *R*_*e*_, so that *θ*_*min*_ becomes close to *π*/2 (*i*.*e*., tangential orientation). *π* If segments are oriented randomly, the average angle should be close to *π* − 1 (calculation shown in SI). We found that the angles of segments located in the middle of chains were between *π*/2 and *π* − 1, indicating that these segments lied predominantly tangential to the surface (**Fig. S**3**c**). While the angles of end segments located in the droplet interior were between *π*/2 and *π* − 1, end segments near the interface exhibited a tendency to form angles larger than *π* − 1 **(Fig. S**3**b)**. However, these segments remained closer to the angle representing isotropic segments, *π* − 1, than perpendicular segments, *π*. These results demonstrate that despite the tendency of chain ends to align perpendicular to the surface, the majority of chain segments remain primarily tangential to the droplet surface.

Based on our investigation of size and orientation of chain segments at different positions in the droplet, we propose a model for homopolymer conformations in condensates as depicted in **Fig.** 4. Compared with the droplet interior, which we found to be consistent with the bulk phase, the distribution of *R*_*g*_ in the interface shifted to smaller values, indicating more compact chain conformations at the droplet interface. Through our characterization of the conformational preferences of segments of different lengths at different positions, we have discovered that segments located at the chains’ ends slightly expand more in the droplet interface compared to the droplet interior, while the middle segments of the chains are more collapsed in the interface region than in the droplet interior. These end segments are enriched at the interface to minimize the entropy loss during LLPS. At the interface, the chain ends exhibit a tendency to align perpendicularly to the interface, but the majority of the chains are still parallel to the surface. We speculate that these conformational changes originate from the change in the local environment surrounding the chains: Compared to the interior of the droplet, the lower monomer concentration at the interface leads to a decrease in the (effective) solvent quality, thus resulting in chain collapse. Similar compact chain conformations at the interface have been reported previously for hydrophobic homopolymers^49–51^.

**Fig. 4.**
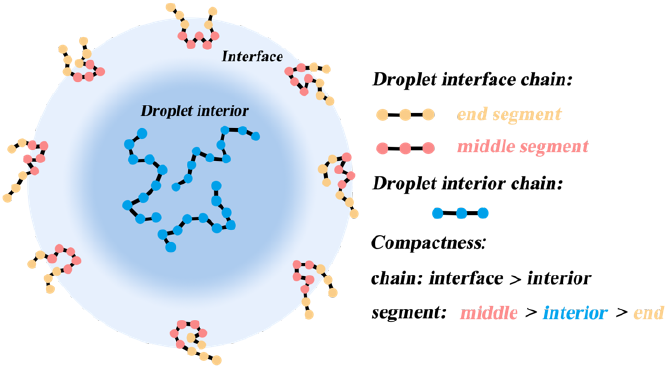
Schematic of homopolymer conformations at the droplet interface. Polymer chains are more compact at the interface than in the droplet interior. Compared to the interior of the droplet, the chain ends are more expanded, while the middle segments of the chains are more collapsed at the interface. The chain ends exhibit a tendency to be perpendicular to the droplet surface, whereas the majority of chains lie predominantly tangential to the droplet surface.

Our findings illustrate the conformational preferences at the condensate interface using a homopolymer model, which can also be observed for naturally occurring IDPs. For FUS LC and LAF-1 RGG (**Fig. S**5), chains are more collapsed at the interface compared with the expanded conformations in the droplet interior. In future work, quantitatively describing the relationship between conformations and interfacial properties will be highly beneficial for our understanding of the biological functions of biomolecular condensates.

## SUPPORTING INFORMATION

The details of the model and simulation, method of average angle calculation, sequences for natural protein (FUS LC and LAF-1 RGG), probability distribution of *R*_*g*_, average *R*_*g*_, *R*_*e*_, angle of different segments at different positions and *R*_*g*_ of natural proteins.

## ACKNOWLEDGMENTS

This material is based on the research supported by the National Institute of General Medical Science of the National Institutes of Health under the grant R01GM136917 and the Welch Foundation under the grant A-2113-20220331. A.N. acknowledges funding by the Deutsche Forschungsgemeinschaft (DFG, German Research Foundation) through Project 470113688. We gratefully acknowledge the computational resources provided by the Texas A&M High Performance Research Computing (HPRC) to complete this work.

## Notes

### Competing Interest Statement

The authors have declared no competing interest.

## REFERENCES

(1) Hyman, A. A.; Weber, C. A.; Jülicher, F. Liquid-Liquid Phase Separation in Biology. Annu. Rev. Cell Dev. Biol. 2014, 30 (1), 39–58. https://doi.org/10.1146/annurev-cellbio-100913-013325.

(2) Banani, S. F.; Lee, H. O.; Hyman, A. A.; Rosen, M. K. Biomolecular Condensates: Organizers of Cellular Biochemistry. Nat. Rev. Mol. Cell Biol. 2017, 18 (5), 285–298. https://doi.org/10.1038/nrm.2017.7.

(3) Alberti, S.; Gladfelter, A.; Mittag, T. Considerations and Challenges in Studying Liquid-Liquid Phase Separation and Biomolecular Condensates. Cell 2019, 176 (3), 419–434. https://doi.org/10.1016/j.cell.2018.12.035.

(4) Dignon, G. L.; Best, R. B.; Mittal, J. Biomolecular Phase Separation: From Molecular Driving Forces to Macroscopic Properties. Annu. Rev. Phys. Chem. 2020, 71 (1), 53–75. https://doi.org/10.1146/annurev-physchem-071819-113553.

(5) Lyon, A. S.; Peeples, W. B.; Rosen, M. K. A Framework for Understanding the Functions of Biomolecular Condensates across Scales. Nat. Rev. Mol. Cell Biol. 2021, 22 (3), 215–235. https://doi.org/10.1038/s41580-020-00303-z.

(6) Uversky, V. N.; Kuznetsova, I. M.; Turoverov, K. K.; Zaslavsky, B. Intrinsically Disordered Proteins as Crucial Constituents of Cellular Aqueous Two Phase Systems and Coacervates. FEBS Lett. 2015, 589 (1), 15–22. https://doi.org/10.1016/j.febslet.2014.11.028.

(7) Brady, J. P.; Farber, P. J.; Sekhar, A.; Lin, Y.-H.; Huang, R.; Bah, A.; Nott, T. J.; Chan, H. S.; Baldwin, A. J.; Forman-Kay, J. D.; Kay, L. E. Structural and Hydrodynamic Properties of an Intrinsically Disordered Region of a Germ Cell-Specific Protein on Phase Separation. Proc. Natl. Acad. Sci. 2017, 114 (39), E8194–E8203. https://doi.org/10.1073/pnas.1706197114.

(8) Martin, E. W.; Mittag, T. Relationship of Sequence and Phase Separation in Protein Low-Complexity Regions. Biochemistry 2018, 57 (17), 2478–2487. https://doi.org/10.1021/acs.biochem.8b00008.

(9) Dignon, G. L.; Zheng, W.; Kim, Y. C.; Best, R. B.; Mittal, J. Sequence Determinants of Protein Phase Behavior from a Coarse-Grained Model. PLOS Comput. Biol. 2018, 14 (1), e1005941. https://doi.org/10.1371/journal.pcbi.1005941.

(10) Schuster, B. S.; Dignon, G. L.; Tang, W. S.; Kelley, F. M.; Ranganath, A. K.; Jahnke, C. N.; Simpkins, A. G.; Regy, R. M.; Hammer, D. A.; Good, M. C.; Mittal, J. Identifying Sequence Perturbations to an Intrinsically Disordered Protein That Determine Its Phase-Separation Behavior. Proc. Natl. Acad. Sci. 2020, 117 (21), 11421–11431. https://doi.org/10.1073/pnas.2000223117.

(11) Mohanty, P.; Kapoor, U.; Sundaravadivelu Devarajan, D.; Phan, T. M.; Rizuan, A.; Mittal, J. Principles Governing the Phase Separation of Multidomain Proteins. Biochemistry 2022, 61 (22), 2443–2455. https://doi.org/10.1021/acs.biochem.2c00210.

(12) Dignon, G. L.; Zheng, W.; Best, R. B.; Kim, Y. C.; Mittal, J. Relation between Single-Molecule Properties and Phase Behavior of Intrinsically Disordered Proteins. Proc. Natl. Acad. Sci. 2018, 115 (40), 9929–9934. https://doi.org/10.1073/pnas.1804177115.

(13) Devarajan, D. S.; Wang, J.; Nikoubashman, A.; Kim, Y. C.; Mittal, J. Sequence-Dependent Material Properties of Biomolecular Condensates. bioRxiv May 12, 2023, p 2023.05.09.540038. https://doi.org/10.1101/2023.05.09.540038.

(14) Brangwynne, C. P.; Tompa, P.; Pappu, R. V. Polymer Physics of Intracellular Phase Transitions. Nat. Phys. 2015, 11 (11), 899–904.

(15) Soranno, A. Physical Basis of the Disorder-Order Transition. Arch. Biochem. Biophys. 2020, 685, 108305. https://doi.org/10.1016/j.abb.2020.108305.

(16) Welles, R. M.; Sojitra, K. A.; Garabedian, M. V.; Xia, B.; Regy, R. M.; Gallagher, E. R.; Mittal, J.; Good, M. C. Determinants of Disordered Protein Co-Assembly Into Discrete Condensed Phases. bioRxiv March 12, 2023, p 2023.03.10.532134. https://doi.org/10.1101/2023.03.10.532134.

(17) Dignon, G. L.; Zheng, W.; Kim, Y. C.; Mittal, J. Temperature-Controlled Liquid–Liquid Phase Separation of Disordered Proteins. ACS Cent. Sci. 2019, 5 (5), 821–830.

(18) Das, R. K.; Pappu, R. V. Conformations of Intrinsically Disordered Proteins Are Influenced by Linear Sequence Distributions of Oppositely Charged Residues. Proc. Natl. Acad. Sci. 2013, 110 (33), 13392–13397. https://doi.org/10.1073/pnas.1304749110.

(19) Sawle, L.; Ghosh, K. A Theoretical Method to Compute Sequence Dependent Configurational Properties in Charged Polymers and Proteins. J. Chem. Phys. 2015, 143 (8), 085101. https://doi.org/10.1063/1.4929391.

(20) Lin, Y.-H.; Forman-Kay, J. D.; Chan, H. S. Theories for Sequence-Dependent Phase Behaviors of Biomolecular Condensates. Biochemistry 2018, 57 (17), 2499–2508. https://doi.org/10.1021/acs.biochem.8b00058.

(21) Devarajan, D. S.; Rekhi, S.; Nikoubashman, A.; Kim, Y. C.; Howard, M. P.; Mittal, J. Effect of Charge Distribution on the Dynamics of Polyampholytic Disordered Proteins. Macromolecules 2022, 55 (20), 8987–8997. https://doi.org/10.1021/acs.macromol.2c01390.

(22) Ghosh, K.; Huihui, J.; Phillips, M.; Haider, A. Rules of Physical Mathematics Govern Intrinsically Disordered Proteins. Annu. Rev. Biophys. 2022, 51, 355–376. https://doi.org/10.1146/annurev-biophys-120221-095357.

(23) Bauer, D. J.; Stelzl, L. S.; Nikoubashman, A. Single-Chain and Condensed-State Behavior of HnRNPA1 from Molecular Simulations. J. Chem. Phys. 2022, 157 (15), 154903. https://doi.org/10.1063/5.0105540.

(24) Rekhi, S.; Sundaravadivelu Devarajan, D.; Howard, M. P.; Kim, Y. C.; Nikoubashman, A.; Mittal, J. Role of Strong Localized vs Weak Distributed Interactions in Disordered Protein Phase Separation. J. Phys. Chem. B 2023, 127 (17), 3829–3838. https://doi.org/10.1021/acs.jpcb.3c00830.

(25) Yu, M.; Heidari, M.; Mikhaleva, S.; Tan, P. S.; Mingu, S.; Ruan, H.; Reinkemeier, C. D.; Obarska-Kosinska, A.; Siggel, M.; Beck, M.; Hummer, G.; Lemke, E. A. Visualizing the Disordered Nuclear Transport Machinery in Situ. Nature 2023, 617 (7959), 162–169. https://doi.org/10.1038/s41586-023-05990-0.

(26) Yuan, X.; W. Hatch, H.; C. Conrad, J.; B. Marciel, A.; C. Palmer J. PH Response of Sequence-Controlled Polyampholyte Brushes. Soft Matter 2023, 19 (23), 4333–4344. https://doi.org/10.1039/D3SM00447C.

(27) Perry, S. L. Phase Separation: Bridging Polymer Physics and Biology. Curr. Opin. Colloid Interface Sci. 2019, 39, 86–97. https://doi.org/10.1016/j.cocis.2019.01.007.

(28) Kelley, F. M.; Favetta, B.; Regy, R. M.; Mittal, J.; Schuster, B. S. Amphiphilic Proteins Coassemble into Multiphasic Condensates and Act as Biomolecular Surfactants. Proc. Natl. Acad. Sci. 2021, 118 (51), e2109967118. https://doi.org/10.1073/pnas.2109967118.

(29) Gouveia, B.; Kim, Y.; Shaevitz, J. W.; Petry, S.; Stone, H. A.; Brangwynne, C. P. Capillary Forces Generated by Biomolecular Condensates. Nature 2022, 609 (7926), 255–264. https://doi.org/10.1038/s41586-022-05138-6.

(30) Böddeker, T. J.; Rosowski, K. A.; Berchtold, D.; Emmanouilidis, L.; Han, Y.; Allain, F. H. T.; Style, R. W.; Pelkmans, L.; Dufresne, E. R. Non-Specific Adhesive Forces between Filaments and Membraneless Organelles. Nat. Phys. 2022, 18 (5), 571–578. https://doi.org/10.1038/s41567-022-01537-8.

(31) Lipiński, W. P.; Visser, B. S.; Robu, I.; Fakhree, M. A. A.; Lindhoud, S.; Claessens, M. M. A. E.; Spruijt, E. Biomolecular Condensates Can Both Accelerate and Suppress Aggregation of α-Synuclein. Sci. Adv. 2022, 8 (48), eabq6495. https://doi.org/10.1126/sciadv.abq6495.

(32) Qin, J.; Priftis, D.; Farina, R.; Perry, S. L.; Leon, L.; Whitmer, J.; Hoffmann, K.; Tirrell, M.; de Pablo, J. J. Interfacial Tension of Polyelectrolyte Complex Coacervate Phases. ACS Macro Lett. 2014, 3 (6), 565–568. https://doi.org/10.1021/mz500190w.

(33) Mitrea, D. M.; Chandra, B.; Ferrolino, M. C.; Gibbs, E. B.; Tolbert, M.; White, M. R.; Kriwacki, R. W. Methods for Physical Characterization of Phase-Separated Bodies and Membrane-Less Organelles. J. Mol. Biol. 2018, 430 (23), 4773–4805. https://doi.org/10.1016/j.jmb.2018.07.006.

(34) Erkamp, N. A.; Farag, M.; Qian, D.; Sneideris, T.; Welsh, T. J.; Ausserwöger, H.; Weitz, D. A.; Pappu, R. V.; Knowles, T. P. J. Adsorption of RNA to Interfaces of Biomolecular Condensates Enables Wetting Transitions. bioRxiv January 13, 2023, p 2023.01.12.523837. https://doi.org/10.1101/2023.01.12.523837.

(35) Dai, Y.; Chamberlayne, C. F.; Messina, M. S.; Chang, C. J.; Zare, R. N.; You, L.; Chilkoti, A. Interface of Biomolecular Condensates Modulates Redox Reactions. Chem 2023, 9 (6), 1594–1609. https://doi.org/10.1016/j.chempr.2023.04.001.

(36) Bratek-Skicki, A.; Van Nerom, M.; Maes, D.; Tompa, P. Biological Colloids: Unique Properties of Membraneless Organelles in the Cell. Adv. Colloid Interface Sci. 2022, 310, 102777. https://doi.org/10.1016/j.cis.2022.102777.

(37) Agudo-Canalejo, J.; Schultz, S. W.; Chino, H.; Migliano, S. M.; Saito, C.; Koyama-Honda, I.; Stenmark, H.; Brech, A.; May, A. I.; Mizushima, N.; Knorr, R. L. Wetting Regulates Autophagy of Phase-Separated Compartments and the Cytosol. Nature 2021, 591 (7848), 142–146. https://doi.org/10.1038/s41586-020-2992-3.

(38) Burgess, D.; Sahin, N. O. Interfacial Rheological and Tension Properties of Protein Films. J. Colloid Interface Sci. 1997, 189 (1), 74–82. https://doi.org/10.1006/jcis.1997.4803.

(39) Besford, Q. A.; Uhlmann, P.; Fery, A. Spatially Resolving Polymer Brush Conformation: Opportunities Ahead. Macromol. Chem. Phys. 2023, 224 (1), 2200180. https://doi.org/10.1002/macp.202200180.

(40) Chang, B.; Zhang, B.; Sun, T. Smart Polymers with Special Wettability. Small 2017, 13 (4), 1503472. https://doi.org/10.1002/smll.201503472.

(41) Tibble, R. W.; Depaix, A.; Kowalska, J.; Jemielity, J.; Gross, J. D. Biomolecular Condensates Amplify MRNA Decapping by Biasing Enzyme Conformation. Nat. Chem. Biol. 2021, 17 (5), 615–623. https://doi.org/10.1038/s41589-021-00774-x.

(42) Farag, M.; Cohen, S. R.; Borcherds, W. M.; Bremer, A.; Mittag, T.; Pappu, R. V. Condensates Formed by Prion-like Low-Complexity Domains Have Small-World Network Structures and Interfaces Defined by Expanded Conformations. Nat. Commun. 2022, 13 (1), 7722. https://doi.org/10.1038/s41467-022-35370-7.

(43) Farag, M.; Borcherds, W. M.; Bremer, A.; Mittag, T.; Pappu, R. V. Phase Separation in Mixtures of Prion-Like Low Complexity Domains Is Driven by the Interplay of Homotypic and Heterotypic Interactions. bioRxiv March 16, 2023, p 2023.03.15.532828. https://doi.org/10.1101/2023.03.15.532828.

(44) Kuo, I. W.; Mundy, C. J.; Eggimann, B. L.; McGrath, M. J.; Siepmann, J. I.; Chen, B.; Vieceli, J.; Tobias, D. J. Structure and Dynamics of the Aqueous Liquid− Vapor Interface: A Comprehensive Particle-Based Simulation Study. J. Phys. Chem. B 2006, 110 (8), 3738–3746.

(45) Fixman, M. Radius of Gyration of Polymer Chains. J. Chem. Phys. 1962, 36 (2), 306–310.

(46) Flory, P. J. Statistical Mechanics of Chain Molecules; Hanser: Munich, 1989.

(47) Midya, J.; Rubinstein, M.; Kumar, S. K.; Nikoubashman, A. Structure of Polymer-Grafted Nanoparticle Melts. ACS Nano 2020, 14 (11), 15505–15516.

(48) Doruker, P.; Mattice, W. L. Simulation of Polyethylene Thin Films on a High Coordination Lattice. Macromolecules 1998, 31 (4), 1418–1426. https://doi.org/10.1021/ma971322z.

(49) Jiang, S.; Luo, C.; Lu, Y. Multilayered Nature in Crystallization of Polymer Droplets Studied by MD Simulations: Orientation and Entanglement. Polymer 2023, 268, 125696. https://doi.org/10.1016/j.polymer.2023.125696.

(50) Kadoya, N.; Arai, N. Size Dependence of Static Polymer Droplet Behavior from Many-Body Dissipative Particle Dynamics Simulation. Phys. Rev. E 2017, 95 (4), 043109. https://doi.org/10.1103/PhysRevE.95.043109.

(51) Berressem, F.; Scherer, C.; Andrienko, D.; Nikoubashman, A. Ultra-Coarse-Graining of Homopolymers in Inhomogeneous Systems. J. Phys. Condens. Matter 2021, 33 (25), 254002. https://doi.org/10.1088/1361-648X/abf6e2.

