## Supporting Information for "Conformational Properties of Polymers at Droplet Interfaces as Model Systems for Disordered Proteins"

### Simulation Details

We have used a coarse-grained model, in which each monomer is represented by a spherical bead that is connected to neighboring beads via a harmonic spring:

$$U_{\mathrm{b}}(r) = \frac{k_{\mathrm{b}}}{2}(r - r_0)^2, \quad (1)$$

where $r$ is the distance between two bonded beads, $k_{\mathrm{b}}$= 20 $\mathrm{kcal}/(\mathrm{mol\AA}^2)$ is the spring constant and $r_0$ = 3.8 Å is the equilibrium length. The van der Waals interactions between nonbonded beads were modeling using a modified Lennard-Jones (LJ) potential[1]

$$U_{\mathrm{vdW}}(r) = \begin{cases} U_{\mathrm{LJ}}(r) + (1-\lambda)\varepsilon, & r \le 2^{1/6}\sigma \\ \lambda U_{\mathrm{LJ}}(r), & otherwise \end{cases}, \quad (2)$$

where $U_{\mathrm{LJ}}$ is the standard LJ potential

$$U_{\mathrm{LJ}}(r) = 4\varepsilon\left[\left(\frac{\sigma}{r}\right)^{12} - \left(\frac{\sigma}{r}\right)^{6}\right], \quad (3)$$

with average hydropathy $\lambda = 0.865$ and bead diameter $\sigma = 6.46$ Å for the homopolymer systems. The interaction strength was fixed to $\varepsilon = 0.2\,\mathrm{kcal/mol}$ in all simulations.

To have good statistics, droplet simulations were performed by placing 5000 chains (50 beads per chain) into a cubic box of edge length 1010 Å with periodic boundary conditions in all directions. We performed Langevin dynamics simulations at temperature $T = 300$K. The friction coefficient was set as $\gamma_i = m_i/t_{\mathrm{damp}}$, where $m_i$ = 163.2 g/mol and $t_{\mathrm{damp}}$ = 1000 fs. Simulations were performed for 2.5 μs with a time step of 10 fs using HOOMD-blue[2] (ver. 2.9.3) with features extended using azplugins[3] (ver. 0.10.2). The simulation trajectories were saved every 1 ns. Analysis was based on the last 1.5 μs.

Bulk system simulations were performed by placing 500 chains into a cubic box at a constant pressure of $P$ = 0 atm for 0.5 μs. After the chains achieved their preferred bulk-phase concentration, the simulation was performed using Langevin dynamics at constant volume for 1 μs. All the simulations were implemented at a fixed temperature of $T$ = 300 K. The analysis was based on the last 0.5 μs. Other simulation settings were the same with droplet simulation.

Single chain simulations were performed by placing one chain into a cubic box of edge length 160 Å. Langevin dynamics simulations were performed for 1 μs at constant temperature $T = 300$ K. The analysis was based on the last 0.5 μs. Other simulation settings were the same with droplet simulation.

Natural protein simulations were performed by placing 1500 chains into a cubic box of edge length 1000 Å with periodic boundary conditions in all directions. Langevin dynamics simulations were performed at temperature $T = 300$K for FUS LC and $T = 260$K for LAF-1 RGG. The hydropathy values ($\lambda_{ij}$) were set according to the HPS-Urry model[4]. The friction coefficient was set as $\gamma_i = m_i / t_{\mathrm{damp}}$, where $m_i$ was the residue mass and $t_{\mathrm{damp}}$ = 1000 fs. Simulations were performed for 1 μs and other simulation settings were the same with droplet forming simulation.

**Natural protein sequences used in this work**

FUS LC (length = 163)

MASNDYTQQATQSYGAYPTQPGQGYSQQSSQPYGQQSYSGYSQSTDTSGYGQSSYSSY
GQSQNTGYGTQSTPQGYGSTGGYGSSQSSQSSYGQQSSYPGYGQQPAPSSTSGSYGSSS
QSSSYGQPQSGSYSQQPSYGGQQQSYGQQQSYNPPQGYGQQNQYNS

LAF-1 RGG (length = 168)

MESNQSNNGGSGNAALNRGGRYVPPHLRGGDGGAAAAASAGGDDRRGGAGGGGYRR
GGGNSGGGGGGGYDRGYNDNRDDRDNRGGSGGYGRDRNYEDRGYNGGGGGGGNRG
YNNNRGGGGGGYNRQDRGDGGSSNFSRGGYNNRDEGSDNRGSGRSYNNDRRDNGGD
G

**Average angle calculation for isotropic distribution of segments**

The average angle of segments distributed without any preferred orientation can be calculated as

$$\theta_{\mathrm{ave}} = \int_{\pi/2}^{\pi} \theta sin\theta \, d\theta = \pi - 1, \quad (4)$$

where $\theta$ is the angle between the segment-to-droplet COM vector and the segment end-to-end vector.

**Supplementary Figures**

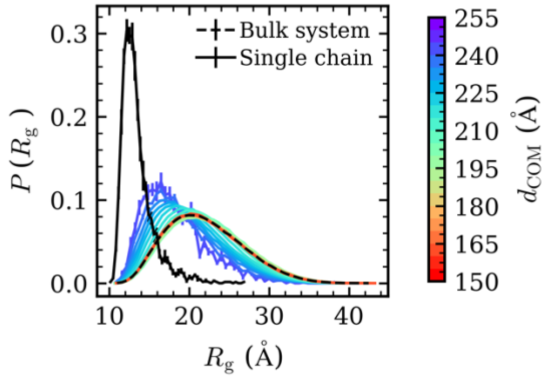


**Fig. S1.** Distribution of radius of gyration ($R_g$). The black dashed line represents the $R_g$distribution in the bulk system. The black solid line represents the $R_g$ distribution of a single chain. The color gradient from red to purple corresponds to the distribution of $R_g$ for chains with $d_{COM}$ ranging from 150 to 255 Å.

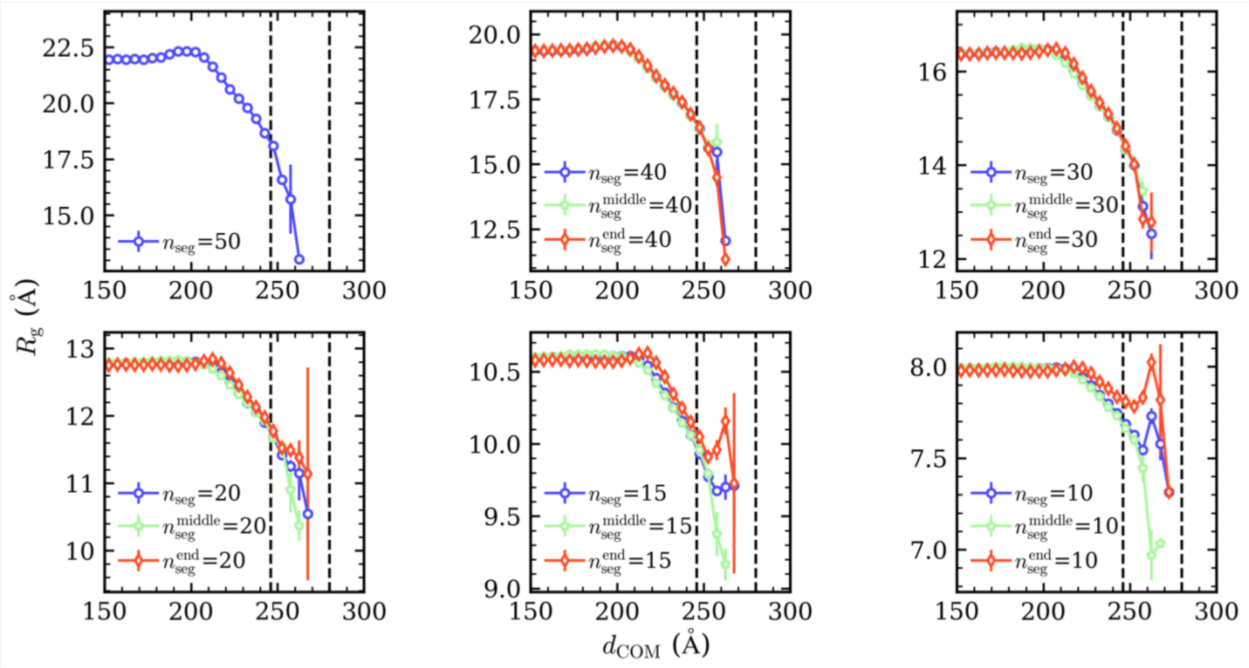


**Fig. S2.** $R_g$ of chains and segments consisting of 40, 30, 20, 15, and 10 monomers as a function of $d_{COM}$. The purple line represents the average $R_g$ for all segments. The green line represents segments located in the middle of the chain, while the red line represents segments located at the chain's ends. The black dashed lines represent the boundaries of the droplet interface.

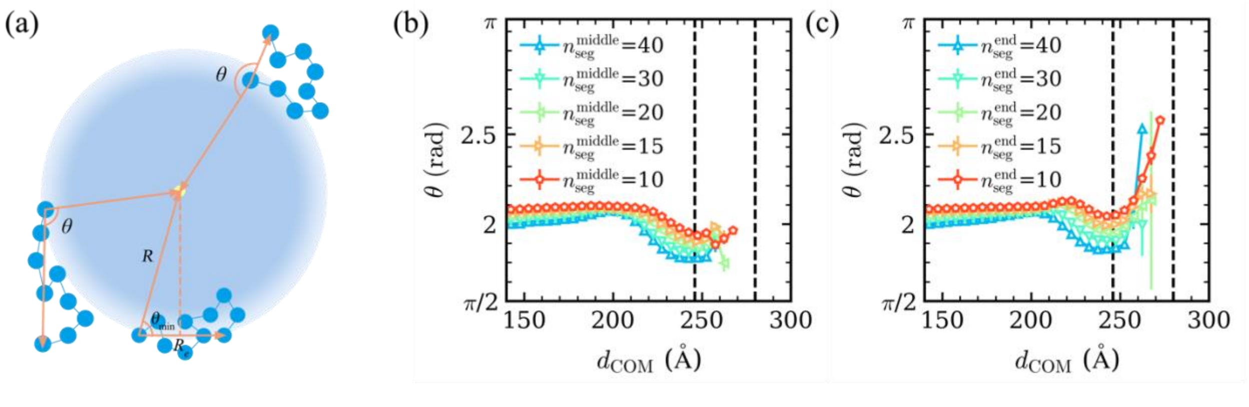


**Fig. S3.** Orientation calculation. (a) Schematic of calculation method for angle ($\theta$) and the minimum of $\theta$ ($\theta_{\min}$). (b) $\theta$ of different length segments located at the middle of a chain with respect to $d_{COM}$. (c) $\theta$ of different length segments located at the ends of a chain with respect to $d_{COM}$.

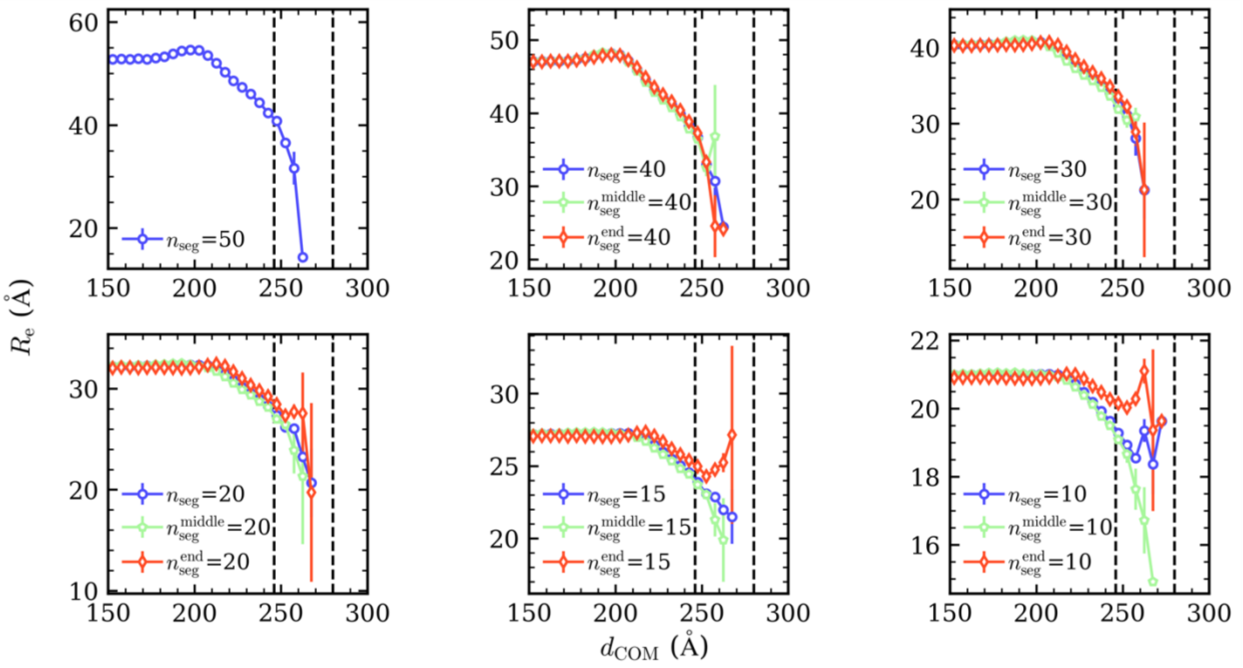


**Fig. S4.** End-to-end distance ($R_e$) of chains and segments consisting of 40, 30, 20, 15, and 10 monomers as a function of $d_{COM}$. The purple line represents the average $R_e$ for all segments. The green line represents segments located in the middle of the chains, and the red line represents segments located at the chain's ends.

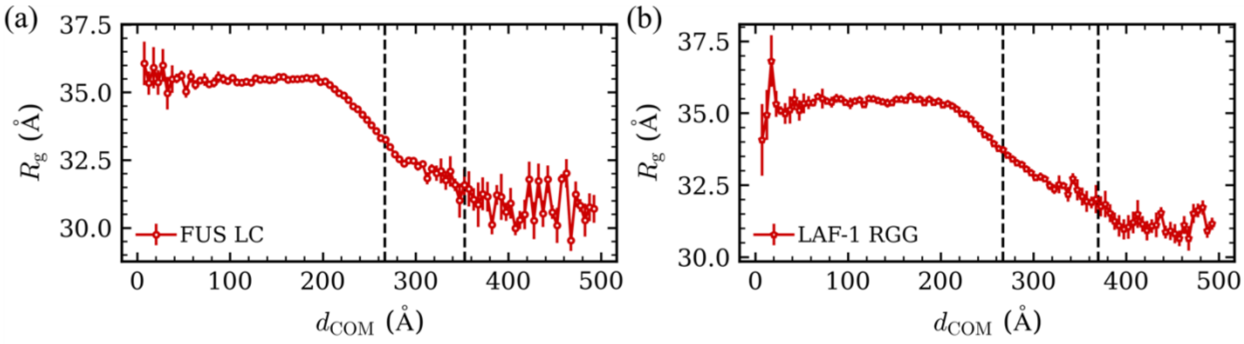


**Fig. S5.** $R_g$ as a function of $d_{COM}$ for (a) FUS LC and (b) LAF-1 RGG sequences. The black dashed lines represent the boundaries of the interface.

### REFERENCES


(1) Ashbaugh, H. S.; Hatch, H. W. Natively Unfolded Protein Stability as a Coil-to-Globule Transition in Charge/Hydropathy Space. *J. Am. Chem. Soc.* **2008**, *130* (29), 9536–9542.

(2) Anderson, J. A.; Glaser, J.; Glotzer, S. C. HOOMD-Blue: A Python Package for High-Performance Molecular Dynamics and Hard Particle Monte Carlo Simulations. *Comput. Mater. Sci.* **2020**, *173*, 109363.

(3) See https://github.com/mphowardlab/azplugins for source code for the software.

(4) Regy, R. M.; Thompson, J.; Kim, Y. C.; Mittal, J. Improved Coarse-Grained Model for Studying Sequence Dependent Phase Separation of Disordered Proteins. *Protein Sci.* **2021**, *30* (7), 1371–1379.
